# Performance analysis of conventional and AI-based variant callers using short and long reads

**DOI:** 10.1101/2023.06.12.544612

**Authors:** Omar Abdelwahab, François Belzile, Davoud Torkamaneh

**Affiliations:** Département de Phytologie, Université Laval, Québec, Canada; Institut de Biologie Intégrative et des Systèmes (IBIS), Université Laval, Québec, Canada; Centre de recherche et d’innovation sur les végétaux (CRIV), Université Laval, Québec, Canada; Institut intelligence et données (IID), Université Laval, Québec, Canada

**Keywords:** genomics, sequencing, variant calling, NGS, artificial intelligence

## Abstract

The accurate detection of variants is essential for genomics-based studies. Currently, there are various tools designed to detect genomic variants, however, it has always been a challenge to decide which tool to use, especially when various major genome projects have chosen to use different tools. Thus far, most of the existing tools were mainly developed to work on short-read data (i.e., Illumina); however, other sequencing technologies (e.g. PacBio, and Oxford Nanopore) have recently shown that they can also be used for variant calling. In addition, with the emergence of artificial intelligence (AI)-based variant calling tools, there is a pressing need to compare these tools in terms of efficiency, accuracy, computational power, and ease of use. In this study, we evaluated the most widely used conventional and AI-based variant calling tools (BCFTools, GATK4, Platypus, DNAscope, and DeepVariant) in terms of accuracy and computational cost using both short-read and long-read data derived from three different sequencing technologies for the same set of samples from the Genome In A Bottle (GIAB) project. The analysis showed that AI-based variant calling tools supersede conventional ones for calling SNVs and INDELs using both long and short reads. In addition, we demonstrate the advantages and drawbacks of each tool while ranking them in each aspect of these comparisons. This study provides best practices for variant calling using AI-based and conventional variant callers with different types of sequencing data.

## Introduction

Genetic variations are defined as any changes in the DNA sequence of individuals within or between species [1]. Next-generation sequencing (NGS) technologies (i.e., second and third generation) have revolutionized the field of genomics by allowing researchers to decode the whole genome of many organisms and genotype very large numbers of genetic variations such as single nucleotide variants (SNVs) and short insertions/deletions (INDELs) [2]. To detect genetic variations, many computational tools, known as variant calling tools or variant callers [3], have also been developed to efficiently identify thousands to millions of variants from sequencing reads aligned against a reference genome. This has been proven to be indispensable in the various areas of genomic research, from agriculture to environment to human health [4–6].

In the last two decades, short reads mainly derived from Illumina sequencing technologies have been the predominant data type used in various variant calling studies [7,8]. Even though short reads provide a high base-level accuracy score, they usually fail to align unambiguously in repetitive regions [9]. While long reads can overcome the challenges posed by repetitive regions, they were not considered suitable for variant calling because of their higher rate of sequencing errors. However, in 2019, Pacific Biosciences (PacBio) introduced a single-molecule real-time (SMRT) sequencing platform that can generate high-fidelity (HiFi) long reads with an average length of 13.5□kilobases (kb) using a Circular Consensus Sequence (CCS) approach. In this approach, a single DNA molecule is circularized, and this template is sequenced multiple times. The resulting consensus provides a sequence with high base-level accuracy (∼99.9%) [10]. Accordingly, HiFi data were used for the detection of genetic variants [10]. In the PrecisionFDA challenge (Truth Challenge V2: Calling Variants from Short and Long Reads in Difficult-to-Map Regions) in 2020, HiFi technology surpassed other sequencing technologies in detecting variants in terms of both precision and recall [11].

Meanwhile, Oxford Nanopore Technologies (ONT) has changed the sequencing paradigm by introducing sequencers that are portable with real-time data delivery and are able to generate ultra-long reads [12]. This technology may look promising for variant calling due to its ability to sequence difficult-to-map regions and read-based phasing, but it has been problematic to achieve a highly accurate analysis because of the error profiles generated by the unique pore-based signal [13]. Nonetheless, recent advances in the development of variant calling tools based on artificial intelligence (AI) (e.g., PEPPER-Margin-DeepVariant) [14] demonstrate that highly accurate variant calling can be achieved from ONT data [14]. Yet, this does not necessarily mean that other variant calling approaches have the ability to detect variants using ONT data.

Over the past few years, besides the advancement of sequencing techniques, many variant calling tools have been developed and used in various genomic projects. For example, the Genome Analysis Toolkit (GATK) [8], developed by the Broad Institute, had been used to detect variants from 180K samples in “The Trans-Omics for Precision Medicine” (TOPMed) program [15]. However, DeepVariant (an AI-based variant caller [16] developed by Google) was selected to detect variants among more than 500K samples by the UK Biobank WES consortium [17], and DRAGEN-GATK [18] was used to genotype more than 1 million samples from the National Institutes of Health’s All of Us Research Program [19]. Despite rapid advances in sequencing technologies and bioinformatics, accurately calling genetic variants from billions of short or error-prone long sequence reads remains challenging. State-of-the-art variant callers use a variety of statistical techniques to distinguish real genetic variants from errors in the reads. However, generalizing these tools to different data types derived from different sequencing technologies has proven difficult. Hence, to date, different variant callers have been used in different NGS-based studies in various species and thus far, it is still challenging to determine which variant calling tool is the best to use. Over the years, various studies have been conducted to compare and evaluate the performance of different variant callers [20–24]. However, all these studies used only short-read sequencing data in their analyses.

In this work, we are addressing the question of whether there is an advantage of a specific variant calling tool over others using a different type of sequencing data (e.g., short vs. long reads). Most of the conventional variant calling tools have been developed and widely used for short-read analysis. However, now with the progress in generating high-quality long reads and the emergence of AI-based variant calling tools, there has been an intriguing question about their potential to supersede conventional ones for calling SNVs and INDELs using both long and short reads. Here, we used three different data types (PacBio HiFi, Illumina, and ONT) for the same set of samples from the Genome in a Bottle Consortium (GIAB) to test five variant callers, two of which are AI-based, in terms of accuracy, time, and computational cost.

## Materials and Methods

### Sequencing data

The sequencing data (Illumina, PacBio HiFi, and ONT) for three samples (HG003, HG006, and HG007) were obtained from the Genome in a Bottle (GIAB) Consortium [25] from the NIST GIAB FTP site: https://ftp-trace.ncbi.nlm.nih.gov/giab/ftp/data/. Only these three samples (out of a total of seven GIAB samples) were used because the other four have been used either to evaluate or train the AI-based variant callers (DeepVariant or DNAscope) [14,16,26].

In summary, Illumina data for HG003, HG006, and HG007 were generated on an Illumina HiSeq2500 and resulted in a lower-than-expected sequence coverage (10.5X, 13.6X, and 12.6X, respectively) due to a large amount of PCR duplicates. The raw FASTQ files were aligned using Sentieon BWA-MEM [27,28], against the GRCh38 reference genome (GCA_000001405.15_GRCh38_no_alt_analysis_set.fna). For the PacBio HiFi data, we downloaded the BAM files directly from the PacBio_CCS_15kb_20kb_chemistry2 directory, as these had been aligned against the same version of the reference genome. In this dataset, the depth of coverage of samples was 42.7X, 40.7X, and 37.6X for HG003, HG005, and HG007, respectively. Finally, ONT data was publicly available only for HG003. Similarly, we downloaded the BAM file directly as it was mapped against the same reference genome. The sequencing coverage of this sample was 77.7X. In this study, we used SAMtools [29] to calculate the coverage of samples using the ‘-a’ option to consider all positions.

### Variant calling

Three conventional variant calling tools (BCFTools [29,30], GATK v4 [8] and Platypus [31]) as well as two AI-based tools (DNAscope [26] and DeepVariant [16]) were used to call variants from the different sets of sequencing data. Variant calling was performed in the same conditions in terms of computational environment. We used default options to filter out low-quality variants. Namely, we followed the recommended parameters for each data type from the BCFTools documentation page and applied only one filter (Quality >=20) after variant calling. For GATK, we utilized the GATK4 pipeline [32] to call variants from Illumina and PacBio data. We started from the variant calling step using the BAM files and performed two rounds of variant calling, where we recalibrated the base quality scores after the first variant calling step to produce recalibrated BAM files for the second round of variant calling. Finally, we used the default filter parameters in the pipeline. For Platypus, variant calling and filtering (default parameters) were conducted following the developers’ recommendations [31]. Platypus was only able to detect variants from Illumina data. To run DNAscope with Illumina data, we started from raw FASTQ files and used the recommended pipeline from Sentieon that includes alignment and duplicate marking. Then, we used the variant calling pipeline that consists of two steps, phasing and a second pass. However, for HiFi data, the whole pipeline is wrapped in a single one-line command (*dnascope_HiFi*.*sh*), as we used the HiFi BAM files directly. As for the ONT data, DNAscope does not have a pipeline for it yet. Finally, for DeepVariant, we followed the documentation on the DeepVariant GitHub repositories to run Illumina, HiFi, and ONT data using the singularity command for each data type.

### Variant calling performance analysis

We identified the common variants between tools and the latest GIAB truth sets v4.2.1 [33] with each data type using the hap.py tool [34]. For evaluation, all the tools were compared in terms of precision (P), recall (R), and F1-score (F1). The equation of each accuracy metric can be represented as 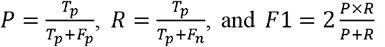, where T_p_, F_p_, and F_n_ stand for true positive, false positive, and false negative, respectively. Here, we presented an average of the performance metrics, however, detailed results for each sample can be found in the supplementary table (Table S1).

### Computational resources and code availability

All the analyses were performed using a Linux system on the Valeria [35] server at Université Laval, QC, Canada. For all variant calling tools, we allowed 16 CPUs and allocated up to 200 GB of RAM to monitor the maximum RAM usage of each tool. The custom code used for the analysis can be accessed on GitHub at https://github.com/Omar-Abd-Elwahab/Variant_Callers.

## Results

### Illumina variant calling performance

#### SNV performance

As can be seen in Figure 1A, DNAscope achieved the highest recall performance (an average of 95.35%). This was ∼2% more than its closest competitor (DeepVariant) and ∼11% more than Platypus, which had the lowest recall performance (84.95%). In terms of precision, DeepVariant (98.95%), Platypus (98.49%), and BCFTools (98.83%) were almost indistinguishable, while DNAscope showed the lowest performance (94.48%). Finally, DeepVariant showed the highest F1-score (96.07%), with a very close performance of BCFTools (95.67%), while Platypus achieved the lowest performance (91.19%).

**Figure 1.**
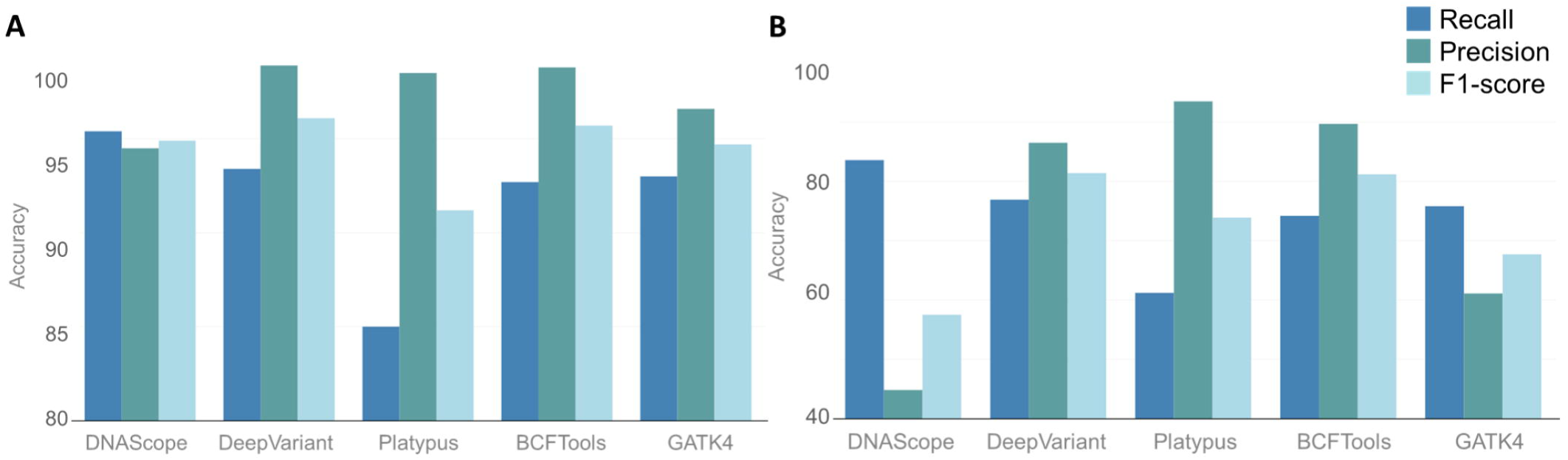
Average accuracy metrics of variants (SNVs (A) and INDELs (B)) called from Illumina data using five different variant callers.

#### INDEL performance

DNAscope achieved the highest recall performance (83.60%) with a difference of ∼6% to its closest competitor (DeepVariant) and ∼22% better than Platypus, which displayed the lowest recall performance (61.17%; Figure 1B). As for precision, Platypus achieved the highest performance (93.53%), while DNAscope had the poorest performance (44.78%), showing a significant difference. The F1-score performance was almost the same for DeepVariant (81.41%) and BCFTools (81.21%), while DNAscope showed the poorest performance (57.53%).

### PacBio HiFi variant calling performance SNV performance

As shown in Figure 2A, DeepVariant and DNAscope demonstrated impressive close-to-perfect performances in all accuracy metrics (>99.9%), although all variant calling tools achieved very high performances (>99%) in all cases. The differences among tools were almost indistinguishable as the differences were less than 1% in precision, recall, or F1-score.

**Figure 2.**
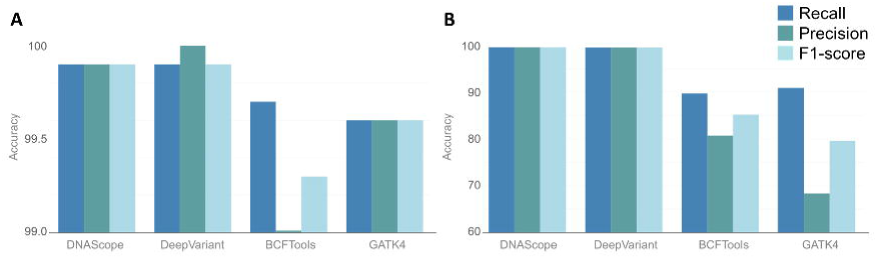
Average accuracy metrics of variants (SNVs (A) and INDELs (B)) called from PacBio HiFi data using four different variant callers.

### INDEL performance

Similarly, DeepVariant and DNAscope achieved the highest performances (>99.5%) in all accuracy metrics. For recall, they were ∼9% more than the closest tool (GATK4) and ∼10% more than BCFTools. In contrast, both BCFTools and GATK4 had a significantly lower precision (<81%). Again, BCFTools and GATK4 saw a significant drop in F1-score, both scoring below 85% (Figure 2B).

### ONT data variant calling performance

To date, BCFTools and DeepVariant (PEPPER-Margin-DeepVariant pipeline) are the only variant callers (out of the five variant callers used in this study) that can handle ONT data. The majority of the SNVs (97.07%) were detected by both DeepVariant and BCFTools. On the other hand, BCFTools failed to detect any INDELs, while DeepVariant had 80.40% in common with the truth sets. Both tools showed a high number of private variants (variants that do not exist in the truth sets) that may be attributed to the quality of ONT sequencing data, resulting in lowering the accuracy metrics even though there is a high number of common variants. In calling SNVs and INDELs, DeepVariant showed a clear advantage over BCFTools in terms of recall, precision, and F1-score (Figure 3).

**Figure 3.**
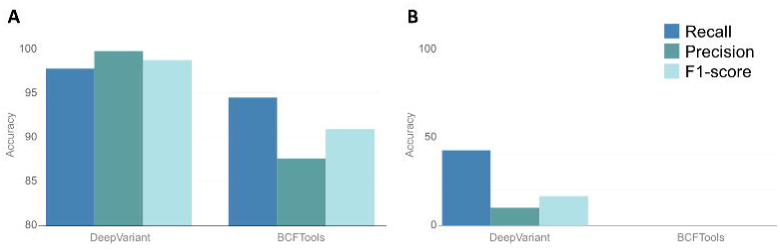
Accuracy metrics of variants (SNVs (A) and INDELs (B)) called from ONT data using BCFTools and DeepVariant with the sample HG003.

### Computational cost of variant calling

As shown in Figure 4, Platypus, DNAscope, and BCFTools proved to be the fastest running tools among the different variant callers (0.34 hours (h), 11.66h, and 7.98h, respectively) for Illumina, PacBio HiFi, and ONT, respectively, whereas GATK4 proved to be the slowest for Illumina and PacBio HiFi requiring 44.19h, and 102.83h, respectively, and DeepVariant was the slowest for ONT data as it required 105.22h.

**Figure 4.**
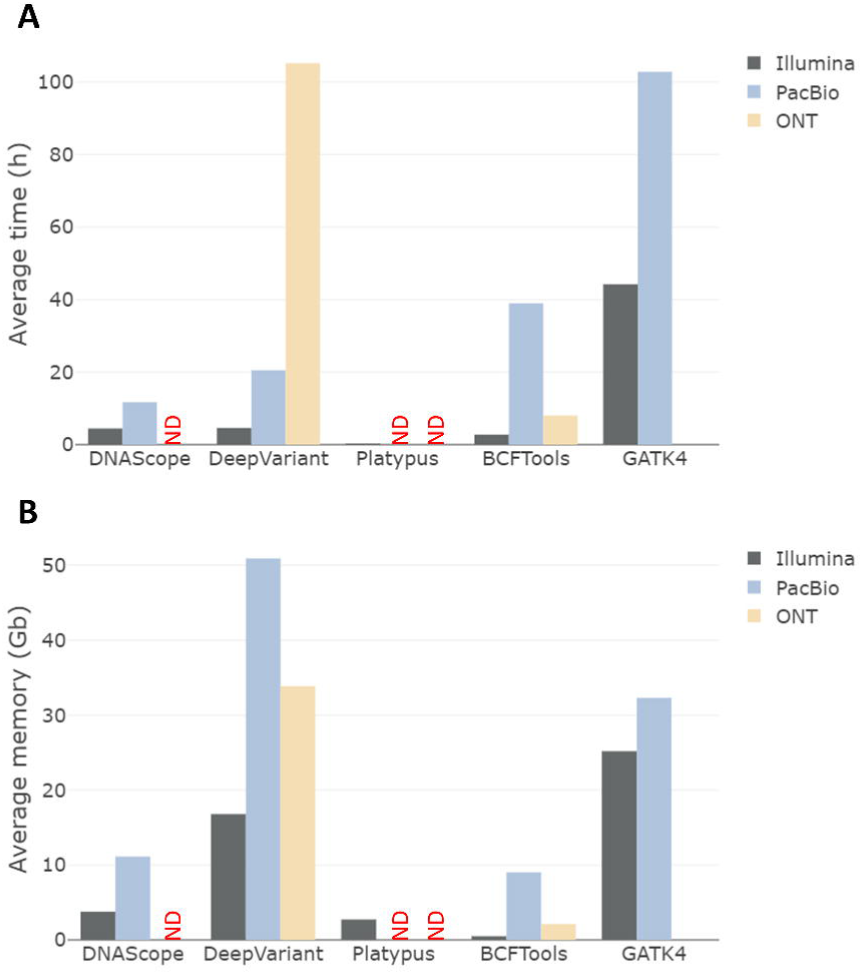
Average computational cost (time (A) and memory (B)) for variant calling using three different data types: Illumina, PacBio HiFi, and ONT. ND: not determined.

In terms of memory required, here also, very large differences were observed. BCFTools proved to be the most memory-efficient tool, requiring 0.49, 9.03, and 2.12 GigaBytes (Gb) to carry out the analyses using Illumina, PacBio HiFi, and ONT data, respectively. GATK4 showed the highest memory usage to process both Illumina and PacBio HiFi data, while DeepVariant was the slowest to process ONT data. However, PacBio and ONT consume much more memory, especially when using DeepVariant, where it reached 50.89 Gb and 33.85 Gb for PacBio and ONT data, respectively.

## Discussion

### Variant calling performance

For years, Illumina has been the benchmark sequencing technology for variant calling despite its difficulty in detecting variants from genomic regions that are considered difficult-to-map by short-read sequencing [36]. However, now with the emergence of long-read sequencing technologies (i.e., PacBio HiFi and ONT), there is a crucial inquiry of whether the conventional variant calling methods will function comparably. Furthermore, there has been a recent emergence of AI-powered variant callers, and researchers are now keen to investigate whether these tools will surpass conventional ones. This study focuses on these concerns by examining various variant callers by utilizing GIAB benchmark data obtained from diverse sequencing technologies [25].

In this study, in alignment with previous comparative studies [2,37–40], a high variant calling accuracy was observed using conventional tools with Illumina data. These studies have proven that conventional variant calling tools generate similar results with a variant concordance of 80-90% on average, where most differences correlate to variants of low coverage or low confidence. As for the AI-based tools, DNAscope proved the most powerful in correctly calling known variants found in the truth sets, but it did have the greatest tendency to call false INDELs using Illumina data resulting in a high-recall-low-precision output. The reason for this could be the insufficient coverage of the Illumina sequencing data. However, on the other hand, DeepVariant provided the most balanced calls combining the best scores for all three metrics considered, as well as the highest F1-score on both SNVs and INDELs. These results are consistent with the original publication of DeepVariant that evaluated DeepVariant’s methodology against multiple conventional tools including GATK and SAMtools and mentioned that DeepVariant demonstrates >50% fewer errors per genome [16]. This superior performance was achieved without entailing a much higher computational cost. Moreover, Olson et al. [11] documented that, with higher coverage (35X), better accuracy metrics can be achieved with DNAscope and DeepVariant, in which the harmonic mean of the F1-score for combined INDELs and SNVs can reach 0.999. Thus, the increasing popularity and use of low-coverage sequencing can pose a challenge for these tools.

PacBio HiFi has been the preferred sequencing technology by many sequencing initiative projects such as the Earth BioGenome Project [41,42], the Vertebrate Genome Project [43], the i5K Initiative [44], and the Ag100Pest Initiative [45]. These projects, among others, have opted for PacBio HiFi technology to produce high-quality reads, making PacBio HiFi the gold standard for generating high-confidence long-reads [46–48]. Although conventional variant callers have been competitive with AI-based tools in identifying variants using Illumina short reads, our study revealed that the AI-based tools exhibited a clear advantage in calling INDELs using PacBio HiFi data, leading to a high-recall-high-precision output. Although there were no significant differences between the tools in calling SNVs using PacBio HiFi data, the AI-based tools showed slightly better performance. Additionally, the AI-based tools outperformed the conventional ones in terms of time and memory efficiency, with DNAscope demonstrating the highest efficiency. Our findings align with the results of the “PrecisionFDA Truth Challenge V2”, where the AI-based tools were the top performers in calling variants in all benchmark regions and difficult-to-map regions from PacBio HiFi data [11]. Previous studies have also shown that using PacBio HiFi data alone could yield equal or better performance to short-read sequencing in all benchmarking regions when calling variants using a single sequencing technology [10,11]. It should be noted that the study utilized a high sequencing coverage (∼40X), therefore it would be valuable to assess the effectiveness and precision of these tools when working with low-coverage data, which is becoming increasingly prevalent.

As for ONT data, previous studies [7,14,49,50] have demonstrated its ability to call genomic variants. In our study, the AI-based variant caller, DeepVariant, showed better results than BCFTools in terms of SNV and INDEL performances using ONT data. However, BCFTools would be a better option in terms of time and memory efficiency when working with ONT data. As documented, it is capable of running on a low to medium-power computer. The results obtained from the variant calling with DeepVariant in this study for SNVs are consistent with the results of recent benchmark studies for ONT data [14,49,50]. However, on the contrary, the INDELs results of this study disagree with the original publication of DeepVariant where the authors have reported higher accuracy. This is probably due to the preprocessing step, in which they used raw ONT reads, carried out the alignment with minimap2 [51], and performed phasing and haplottaging [16]. However, here, we performed the standard procedures by running the default code on the acquired BAM files from GIAB directly for generalization between tools, and the possibility of data unavailability in some variant calling projects. A recent AI-based variant caller specifically designed for ONT data, CLAIR3 [52], has shown that it can achieve similar results to DeepVariant in calling variants. However, the “PrecisionFDA Truth Challenge V2” has mentioned DeepVariant as a top performer in calling variants from ONT data, especially from difficult-to-map regions [11]. Moreover, another study has claimed that highly accurate variants (94.25% F1-score) can be called with lower coverage in ONT data [49]. This suggests that ONT data can be used for reliable variant calling, but there is still room for improvement in the accuracy and efficiency of the tools used for this purpose.

Overall, PacBio HiFi and ONT data (long reads) have the ability to compete with Illumina (short reads) in calling genomic variations. Furthermore, utilizing AI-based variant calling tools with both short and long reads can achieve very high accuracy metrics for calling both SNVs and INDELs. Namely, DeepVariant has overall better performance with all data types even with comparatively lower coverage as in the Illumina case. Recent studies have shed light on the fact that combining sequencing technologies produce better accuracy than any separate sequencing technology [42]. This encourages the production of more AI-based tools that can call variants from multiple technologies at the same time to achieve better results.

### User experience

Although it was very smooth to set up all the tools, running them did not give the same experience. For the AI-based tools, the documentation was extremely clear and helpful. Especially when utilizing DNAscope with PacBio HiFi data, all the steps are compacted in a single command line. As a commercial tool, the DNAscope support team was very accessible and easy to reach. As for DeepVariant, it is always a singularity command that can perform all the processes of variant calling no matter the data type. Moreover, both tools do not require filtering variants or setting thresholds manually to refine the results.

On the other hand, conventional tools led to a different experience. Both BCFTools and Platypus are very easy to handle with very clear documentation. However, Platypus is still a Python 2-dependent tool, only works on short reads, and has not been updated since 2014. In contrast, BCFTools has been improved and updated regularly over the years. Platypus includes default values to filter variants, while for BCFTools and GATK4 all the filters need to be set manually. Running GATK requires an in-depth understanding of all the steps and parameters to set manually. Although this gives the user more control over the filtering process, setting thresholds for the filters might be an exhausting and time-consuming process.

The AI-based variant calling tools have an advantage in user time performance due to their automatic filtration feature which makes them less time-consuming overall. The time performance was only calculated for the variant calling step without taking into consideration setting thresholds, filtering, or any other pre-/processing steps.

## Limitations

Although PacBio HiFi data achieved the best results, the high coverage (∼40X) might be the reason behind this advantage over Illumina data (∼12X). According to [14], PEPPER-Margin-DeepVariant can achieve better results in calling INDELs when following specific procedures that include performing phasing and haplotagging. Moreover, we compared the capability of each tool in its default form to mimic the conditions of regular users.

## Conclusion

Currently, the long-read data show the potential to become the new standard for variant calling and genotyping. PacBio HiFi introduces low error rates with high base calling quality while having an edge in detecting repetitive regions that are difficult to handle with short-read technologies. Utilizing PacBio HiFi data is now leading to near-optimal SNV and INDEL performance competing with short-read technologies. The long reads are also the optimal technique to detect structural variants allowing now to identify all types of genetic variations with a single sequencing experiment. The only drawback of long-read technologies, which is making it behind short-read technologies, is the cost where Illumina still has the edge of being the cheapest in the market.

Combining with long reads, AI-based tools have demonstrated a clear advantage over conventional tools in calling variants, which paves the road and makes it a starting point for a new era of AI tools in the genomics field. As noticed in this article, AI-based tools do not perform in the same time frame, which might be because of the engineering design of each tool. Being said, this concludes that there is still room for improvements in AI-based tools, where they can even give better performances that might reach the gold standards in the future, achieving less computational cost and more efficacy.

## Key points

- We did performance analyses of conventional and AI-based variant calling pipelines to test their abilities to accurately call genomic variations such as SNPs and small INDELs from various NGS data such as Illumina, PacBio HiFi, and ONT.
- The analyses show the differences among variant calling pipelines in terms of accuracy metrics and computational cost.
- We discuss the best practices for various variant calling pipelines while reporting the user experience.

## Acknowledgments

The authors wish to thank Génome Québec, Genome Canada, the government of Canada, the Ministère de l’Économie et de l’Innovation du Québec, the Canadian Field Crop Research Alliance, Semences Prograin Inc., Sollio Agriculture, Grain Farmers of Ontario, Barley Council of Canada, and Université Laval. The authors wish also to thank GIAB for providing the reference variants of the samples. Moreover, we thank Mr. Don Freed at Sentieon for his continuous fast responses and guidance regarding the best practices of the usage of DNAscope.

## Authors’ Contributions

Conceptualization, O.A. and D.T.; data curation and formal analysis, O.A.; resources, D.T.; writing, review and editing, O.A., D.T., and F.B.; supervision, D.T.; project administration, D.T.; funding acquisition, D.T., and F.B. All authors have read and agreed to the published version of the manuscript.

## Author Biographies

**Omar Abdelwahab** is a Ph.D. student at Laval University. He did his bachelor’s degree in genomics and computational biology from the Biomedical Sciences department at Zewail University of Science and Technology. His research mainly focuses on developing AI-based genomic pipelines and analyzing genomic methods and data.

**Francois Belzile** is a professor (Laval University) and a world expert in plant genomics. Over the past few years, he has been leading several large-scale genomics projects funded by Genome Canada.

**Davoud Torkamaneh** is a professor (Laval University) and expert in the field of computational biology, with a strong track record of translating genomics and big data to real-world applications. He has contributed significantly to the development of bioinformatics tools that are commonly used by various research groups.

