## supplementary table (Table S1) for "Performance analysis of conventional and AI-based variant callers using short and long reads"

**Table S1.** Detailed results for performance analysis of calling variants from three samples using five variant calling tools with different sequencing technologies (Illuina, Pacbio HiFi, and  ONT) in terms of accuracy metrics (precision, recall and F1-score), running time, and memory**.**

| | | | | | | | | AI-based | | | | | | | | | | Conventional |
|---|---|---|---|---|---|---|---|---|---|---|---|---|---|---|---|---|---|---|
| | | | | | DNAscope | | | DeepVariant | | | Platypus | | | BCFTools | | | | GATK4 |
| | | | Illumina | PacBio HiFi | ONT | Illumina | PacBio HiFi | ONT | Illumina | PacBio HiFi | ONT | Illumina | PacBio HiFi | ONT | | Illumina | PacBio HiFi | ONT |
| | | Precsion | 0.5878 | 0.9955 | | 0.8449 | 0.9949 | 0.1035 | 0.9257 | | | 0.8775 | 0.8300 | 0.0000 | | 0.5842 | 0.7422 | |
| | INDEL | Recall | 0.8071 | 0.9954 | | 0.7317 | 0.9949 | 0.4232 | 0.5471 | | | 0.6925 | 0.8829 | 0.0004 | | 0.7110 | 0.8969 | |
| | | F1score | 0.6802 | 0.9955 | | 0.7842 | 0.9949 | 0.1663 | 0.6878 | | | 0.7741 | 0.8557 | 0.0000 | | 0.6414 | 0.8123 | |
| | | Precsion | 0.9620 | 0.9993 | | 0.9867 | 0.9995 | 0.9968 | 0.9817 | | | 0.9861 | 0.9894 | 0.8759 | | 0.9600 | 0.9955 | |
| HG003 | SNP | Recall | 0.9418 | 0.9994 | | 0.9213 | 0.9992 | 0.9769 | 0.8018 | | | 0.9032 | 0.9964 | 0.9445 | | 0.9129 | 0.9960 | |
| | | F1score | 0.9518 | 0.9993 | | 0.9528 | 0.9994 | 0.9868 | 0.8827 | | | 0.9428 | 0.9929 | 0.9089 | | 0.9358 | 0.9958 | |
| | | Time (hours) | 3.4000 | 11.8333 | | 5.1833 | 35.6833 | 105.2167 | 0.2580 | | | 2.5500 | 36.7500 | 7.9833 | | 36.6000 | 100.7667 | |
| | | Memory (GB) | 3.3100 | 10.7100 | | 17.3300 | 51.0800 | 33.8500 | 2.8500 | | | 0.4938 | 6.9500 | 2.1200 | Round 1 | 20.6500 | 42.8000 | |
| | | | | | | | | | | | | | | | Round 2 | 27.5200 | 29.2200 | |
| | | Precsion | 0.3588 | 0.9969 | | 0.8770 | 0.9965 | | 0.9410 | | | 0.9076 | 0.8553 | | | 0.6127 | 0.7446 | |
| | INDEL | Recall | 0.8557 | 0.9968 | | 0.7940 | 0.9969 | | 0.6506 | | | 0.7775 | 0.9125 | | | 0.7908 | 0.9233 | |
| | | F1score | 0.5056 | 0.9968 | | 0.8334 | 0.9967 | | 0.7693 | | | 0.8375 | 0.8830 | | | 0.6905 | 0.8244 | |
| | | Precsion | 0.9328 | 0.9994 | | 0.9917 | 0.9996 | | 0.9870 | | | 0.9898 | 0.9903 | | | 0.9675 | 0.9957 | |
| HG006 | SNP | Recall | 0.9623 | 0.9995 | N/A | 0.9419 | 0.9994 | | 0.8799 | N/A | N/A | 0.9452 | 0.9974 | | | 0.9423 | 0.9961 | N/A |
| | | F1score | 0.9473 | 0.9994 | | 0.9662 | 0.9995 | | 0.9304 | | | 0.9670 | 0.9939 | | | 0.9547 | 0.9959 | |
| | | Time (hours) | 5.2333 | 10.7333 | | 4.1667 | 24.0833 | | 0.3833 | | | 2.9500 | 40.3333 | | | 50.1833 | 96.8500 | |
| | | Memory (GB) | 4.1500 | 13.3000 | | 16.4600 | 53.1500 | | 2.6500 | | | 0.4938 | 7.9200 | | Round 1 | 24.9000 | 23.9000 | |
| | | | | | | | | N/A | | | | | | N/A | Round 2 | 23.8700 | 28.1100 | |
| | | Precsion | 0.3968 | 0.9953 | | 0.8728 | 0.9924 | | 0.9394 | | | 0.9050 | 0.7360 | | | 0.6376 | 0.5625 | |
| | INDEL | Recall | 0.8452 | 0.9950 | | 0.7814 | 0.9933 | | 0.6374 | | | 0.7576 | 0.8941 | | | 0.7718 | 0.9074 | |
| | | F1score | 0.5400 | 0.9952 | | 0.8246 | 0.9928 | | 0.7595 | | | 0.8247 | 0.8074 | | | 0.6983 | 0.6945 | |
| | | Precsion | 0.9395 | 0.9993 | | 0.9900 | 0.9995 | | 0.9861 | | | 0.9891 | 0.9898 | | | 0.9692 | 0.9961 | |
| HG007 | SNP | Recall | 0.9565 | 0.9993 | | 0.9375 | 0.9992 | | 0.8668 | | | 0.9334 | 0.9970 | | | 0.9335 | 0.9957 | |
| | | F1score | 0.9479 | 0.9993 | | 0.9631 | 0.9993 | | 0.9226 | | | 0.9604 | 0.9934 | | | 0.9511 | 0.9959 | |
| | | Time (hours) | 4.5500 | 12.4167 | | 4.3000 | 19.9500 | | 0.3833 | | | 2.8000 | 39.8500 | | | 45.7833 | 110.8667 | |
| | | Memory (GB) | 3.8500 | 9.4000 | | 16.5600 | 48.4500 | | 2.6700 | | | 0.4943 | 12.2300 | | Round1 | 24.9900 | 36.3500 | |
| | | | | | | | | | | | | | | | Round2 | 29.2400 | 33.3600 | |
